# Alcohol induces concentration-dependent transcriptomic changes in oligodendrocytes

**DOI:** 10.1101/2023.09.22.559075

**Authors:** Sam A. Bazzi, Cole Maguire, R. Dayne Mayfield, Esther Melamed

**Affiliations:** Department of Neurology, Dell Medical School, The University of Texas at Austin, Austin, TX, USA; Department of Neuroscience, The University of Texas at Austin, Austin, TX, USA

## Abstract

Oligodendrocytes are a key cell type within the central nervous system (CNS) that generate the myelin sheath covering axons, enabling fast propagation of neuronal signals. Alcohol consumption is known to affect oligodendrocytes and white matter in the CNS. However, most studies have focused on fetal alcohol spectrum disorder and severe alcohol use disorder. Additionally, the impact of alcohol dosage on oligodendrocytes has not been previously investigated. In this study, we evaluated transcriptomic changes in C57BL6/J cultured mature oligodendrocytes following exposure to moderate and high concentrations of alcohol. We found that high concentrations of alcohol elicited gene expression changes across a wide range of biological pathways, including myelination, protein translation, integrin signaling, cell cycle regulation, and inflammation. Further, our results demonstrate that transcriptomic changes are indeed dependent on alcohol concentration, with moderate and high concentrations of alcohol provoking distinct gene expression profiles. In conclusion, our study demonstrates that alcohol-induced transcriptomic changes in oligodendrocytes are concentration-dependent and may have critical downstream impacts on myelin production. Targeting alcohol-induced changes in cell cycle regulation, integrin signaling, inflammation, or protein translation regulation may uncover mechanisms for modulating myelin production or inhibition. Furthermore, gaining a deeper understanding of alcohol’s effects on oligodendrocyte demyelination and remyelination could help uncover therapeutic pathways that can be utilized independent of alcohol to aid in remyelinating drug design.

## Introduction

Oligodendrocytes (OLs) are glial cells within the central nervous system (CNS) that produce myelin, which coats neuronal axons and enhances fast propagation of neuronal signals. Production of myelin is a metabolically expensive process, making OLs particularly sensitive to environmental insults (Wilhelm & Guizzetti, 2016).

One of the important environmental factors that can impact oligodendrocytes is alcohol, though its effects have been mainly studied in fetal alcohol spectrum disorder (FASD) and in severe alcohol use disorder (AUD). In FASD, alcohol exposure leads to severe abnormalities throughout the brain, but particularly in white matter-rich regions such as the corpus callosum (Sowell et al., 2008). Experiments in animal models of FASD have demonstrated increased OL precursor cell (OPC) apoptosis and reduced OL differentiation (Darbinian et al., 2021). In turn, AUD has been associated with microstructural white matter damage in multiple MRI brain imaging studies which assess myelination status (Pfefferbaum et al., 2014; Chumin, et al., 2019). Additionally, post-mortem studies on human brains from AUD patients have revealed a broad decrease in expression of myelination pathway genes (Lewohl et al., 2000). Alcohol toxicity is also of particular concern in autoimmune demyelinating disorders such as multiple sclerosis (MS).

Nonetheless, there have been few studies on the effects of alcohol on mature oligodendrocytes (mOLs) compared to OPCs. In addition, few studies have focused on the dose-dependent impact of alcohol on mOL phenotypes. These are important questions, as many demyelinating neurological disorders have not been associated with early life exposure to alcohol yet may still be affected by alcohol during the onset of demyelinating disease later in life (Diaz-Cruz, et al., 2017; Andersen, et al., 2018; Ivashynka, et al., 2019). Further, levels of alcohol consumption vary greatly across the population, raising the importance of understanding how moderate versus high levels of alcohol may impact mOL function.

The goal of our study was to evaluate mOL gene expression changes in response to moderate and high concentrations of alcohol in a well-controlled *in vitro* system. The transcriptome is important for identifying molecular changes in response to environmental toxins, and alcohol in particular has been known to impact myelin genes in AUD (Costin and Miles, 2015). Since individual alcohol consumption varies across the population, it is important to understand the impact of both moderate and high levels of consumption on the mOL transcriptome. Furthermore, understanding how different concentrations of alcohol impact the mOL transcriptome may have ramifications for advising patients with demyelinating disorders on alcohol consumption. Our results demonstrate that transcriptomic changes are indeed dependent on alcohol concentration, with moderate and high concentrations of alcohol provoking distinct gene expression profiles.

## Methods

### Tissue collection and OL isolation

All animal work was approved by the University of Texas at Austin (UT Austin) Institutional Animal Care and Use Committee. C57BL6/J mice (post-natal days 5-7), bred in our UT Austin colony, were euthanized by rapid decapitation. Protocol for tissue processing and OL isolation was adapted from Flores-Obando, Freidin, & Abrams, 2018 (Figure 1A). Briefly, the brain was removed, and the cortices pried away. Isolated cortices were digested with papain solution and a series of washes along with mechanical disruption to achieve a single cell suspension. After straining through fine mesh filters, the suspension was counted and incubated with Anti-O4 MicroBeads at a concentration of 10 uL of beads per 1×10^7^ cells (Miltenyi Biotec). After incubation, the suspension was washed and passed through an LS Column placed inside a magnetic separator. The column was washed four times before removing the column from the magnet and eluting the O4-labeled cells. O4-labeled cells were then washed and resuspended at 5 x 10^5^ cells/100 uL of media. 100 uL of media was seeded onto Poly-D-Lysine/Laminin coated coverslips placed inside a 24-well plate. The plate was incubated for an hour inside a cell culture incubator (37° C/5% CO_2_) for 45 minutes to allow cells to adhere to the coverslip before flooding the well with 500 uL of media. After 24 hours, the media was replaced with differentiation medium containing differentiation factor triiodothyronine (T3). Differentiation continued for three additional days before alcohol exposure.

**Figure 1.**
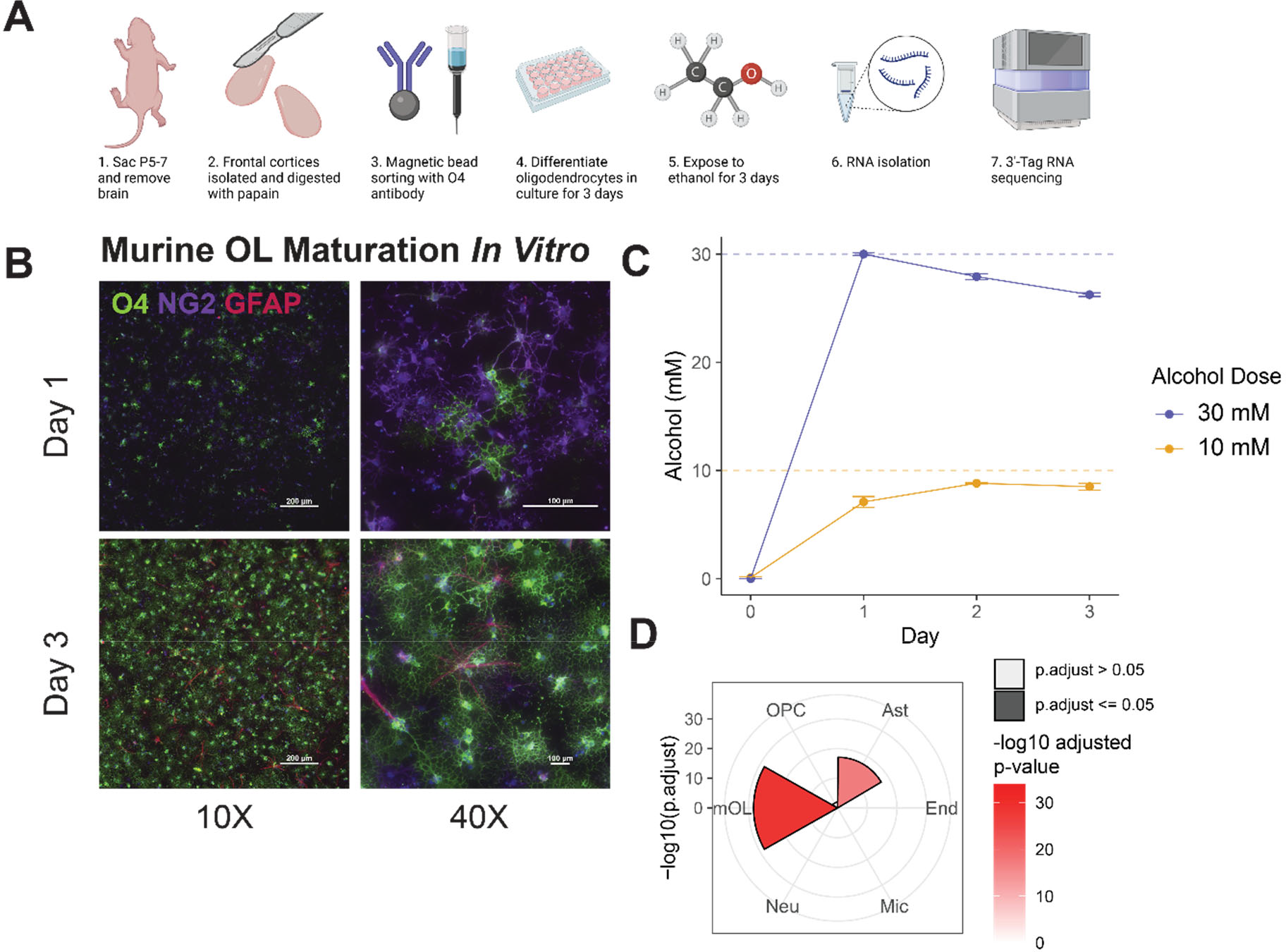
Experiment outline and culture validation. **A)** xsExperimental outline of OL culture, dosing paradigm and sequencing. **B)** Immunocytochemistry of OL marker O4, glial marker NG2, and astrocyte marker GFAP. Over three days, NG2 expression decreases as O4 and GFAP expression increase. Culture is ∼90% pure for OLs. **C)** Alcohol concentration measured over three days for 10 mM and 30 mM dosing. **D)** Cell type enrichment analysis confirmed enrichment of mOL marker genes in the culture.

### Immunofluorescence staining

OL differentiation and astrocyte contamination were monitored by immunofluorescence on Day 1 and Day 3 *in vitro*. NG2 was utilized as an early glial marker, O4 as an OL marker, and GFAP as an astrocyte marker. Coverslips were stained with O4 and NG2 primary antibodies for 45 minutes inside a cell culture incubator (37° C/5% CO_2_). After washing with 0.05% Tween-20/PBS solution, cells were fixed with 4% paraformaldehyde for 10 minutes at room temperature. Coverslips were washed again with Tween solution three times before incubating with secondary antibodies (1:400) diluted in medium for 1 hour at room temperature in the dark. After washing off the secondary antibody, cells were permeabilized with perm/block buffer for internal staining of GFAP. GFAP primary was diluted 1:100 in perm/block buffer and incubated for 1 hour at room temperature. Following this washing step, secondary antibody was added (1:400 in medium) and allowed to incubate for 1 hour at room temperature in the dark. Finally, the coverslips were washed and mounted with DAPI-containing mounting medium onto microscope slides. Coverslips were cured overnight in the dark at room temperature before imaging on a Nikon Eclipse Ni-E upright microscope.

### Alcohol exposure

After three days of differentiating *in vitro*, coverslips with OLs were exposed to alcohol. To mitigate the effects of evaporation on alcohol concentrations, the culture plates were placed in a polystyrene box along with a beaker containing alcohol at a concentration double that of the target concentration in the medium. The polystyrene box was then placed inside a cell culture incubator (37° C/5% CO_2_). This “compensation” method provides a much slower decrease in the concentration of alcohol contained in the medium by providing a steady source of alcohol vapor in the air, along with a smaller microenvironment within the incubator due to the polystyrene box. Alcohol concentration in the medium was sampled using an Analox AM1 machine to confirm that the target concentration was reached and sustained over three days of exposure.

### RNA Isolation and 3’-Tag Sequencing

After three days of alcohol exposure (or no exposure for controls), cultured OLs were lysed for RNA extraction using the Qiagen RNEasy Mini kit according to manufacturer’s instruction. Isolated RNA was quantified using the Qubit 4 with the Qubit RNA BR Assay Kit. RNA was spot checked with a bioanalyzer to confirm high RNA integrity (Supplemental Table 1).

3’ Tag Sequencing of the extracted RNA (using <150 ng) was performed using a NovaSeq SR100 S1 by the Genomic Sequencing and Analysis Facility at UT Austin, Center for Biomedical Research Support (RRID# SCR_021713). 3’ TagSeq was performed by the University of Texas Genomic Sequencing and Analysis Facility, based on the protocols from Lohman BK et al. (2016) and Meyer E et al. (2011). Libraries were quantified using the Quant-it PicoGreen dsDNA assay (ThermoFisher) and pooled equally for subsequent size selection at 350-550bp on a 2% gel using the Blue Pippin (Sage Science). The final pools were checked for size and quality with the Bioanalyzer High Sensitivity DNA Kit (Agilent) and their concentrations were measured using the KAPA SYBR Fast qPCR kit (Roche). Samples were then sequenced on the NovaSeq 6000 (Illumina) instrument with single-end, 100-bp reads. Computational analyses were performed using the Biomedical Research Computing Facility at UT Austin, Center for Biomedical Research Support (RRID#: SCR_021979). Reads were trimmed and filtered with cutadapt (v 1.18) before mapping to the GRCm39 cDNA ensembl mouse transcriptome using STAR (v 020201). Total reads passing filtering was >90% for each sample with >87% of these uniquely mapped (Supplemental Table 1).

Read counts were normalized and filtered for reads with a total of greater than or equal to 10 reads across all samples. Differential expression was calculated using the package DESeq2 (v 1.34.0) defining differentially expressed genes (DEGs) by Benjamini-Hochberg corrected p-values (p.adj) <= 0.05. Log fold change was shrunk using apeglm (Zhu, Ibrahim & Love, 2018), and for 30 mM v 0 mM and 10 mM v 0 mM comparisons, alcohol exposed groups were compared only to 0 mM controls in their batch and sequencing run. Principal component analysis was performed on the top 10,000 most variable genes, using DESeq2 variance stabilized counts normalized to mean expression values of the 0 mM controls on the same batch and sequencing run. Gene Set Enrichment Analysis (GSEA) was conducted on the apeglm shrunk log fold change using the R package ClusterProfiler (v4.2.2) to calculate Kyoto Encyclopedia of Genes and Genomes (KEGG), Reactome, Gene Ontology (GO), Hallmark, and C3 gene set enrichment. To evaluate enrichment for specific cell type marker genes, the mean normalized variance stabilized counts from DESeq2 of all samples were used with GSEA using a gene set of defined CNS cell type marker genes from McKenzie, et al., 2018. For oligodendrocyte cell type subpopulation enrichment, marker genes from Pandey et al., 2022 were used with mean normalized variance stabilized counts of all samples. For calculating percent change of gene expression for 10 mM and 30 mM alcohol exposure, only control samples on the same batch and sequencing run were used. For the ranked-ranked hypergeometric testing and concentration dependent analyses between 10 mM and 30 mM, gene expression was normalized to their batch and sequencing run’s accompanying 0 mM controls.

### Data Availability

Raw and processed 3’ tag-seq data are accessible in Gene Expression Omnibus under accession code GSE239691.

## Results

Tag-sequencing was conducted in two batches, with batch 1 containing 30 mM and 0 mM exposed samples and batch 2 containing 10 mM and 0 mM exposed samples (see methods for discussion on batch correction). Tag-sequencing reads were mapped to a total of 38,979 genes, with a total of 34,851 genes for batch 1 and 36,613 genes for batch 2 (meaning 2,366 genes were mapped only in batch 1 and 4,128 genes were mapped only in batch 2). From these, genes were filtered to those that had at least 10 counts across all samples resulting in a total of 26,992 genes used for analysis (with 23,218 when using only batch 1 samples and 24,386 genes when using only samples in batch 2).

### In vitro oligodendrocyte differentiation and alcohol exposure

OLs were enriched to a purity of approximately 88% as confirmed by immunocytochemistry (Figure 1B, x=0.880 +/-0.0265, n=3). From Day 1 to Day 3, early glial marker neuron-glial antigen (NG2) expression decreased as OL marker O4 expression increased, with low glial fibrillary acidic protein (GFAP) presence indicating 12% of cells were astrocytes.

Alcohol concentration in cell culture medium was measured over three days (Figure 1C). Concentrations of 10mM and 30mM alcohol were chosen for exposure, as 10mM (46.1 mg/dL BAC) corresponds with moderately intoxicated drinking, whereas 30mM (138.3 mg/dL BAC) corresponds with heavy intoxication in humans (Dasgupta, 2017). In the high concentration condition (30mM group), alcohol peaked at 30mM after the first day and dissipated to 27 mM by day 3. In the moderate concentration condition (10mM group), alcohol peaked at 9mM by day 2 and this concentration was maintained to day 3 (Figure 1C).

Cell type enrichment analysis via 3’-Tag sequencing confirmed that mOLs made up the majority of cell types in the culture (Figure 1D), with high expression of mOL-associated genes proteolipid protein 1 *(PLP1)*, myelin basic protein *(MBP)*, and myelin associated oligodendrocyte basic protein *(MOBP)*. Similar to our immunocytochemistry study, we observed a small population of astrocytes in culture defined by gene expression of apolipoprotein E *(APOE), GFAP*, and peroxiredoxin 6 *(PRDX6)* (Supplemental Table 2).

CNS gene markers for various brain cell types were identified from a published data set of hundreds of genes associated with distinct CNS lineages (McKenzie et al., 2018). Based on this dataset, we did not observe significant enrichment in oligodendrocyte progenitor cell (OPC) markers, suggesting successful differentiation of OPCs into mature OLs.

In addition, using marker genes of six known OL subpopulations, (Marques et al., 2016; Pandey et al., 2022) we did not find significant overenrichment of markers belonging to specific subpopulations of mOLs (Supplemental Table 2), suggesting that our culture captured mixed mOL phenotypes. Furthermore, subpopulation marker genes were not significantly different in either the moderate or high concentration alcohol groups, suggesting that alcohol exposure did not affect the relative proportion of the six mOL phenotypes (Supplemental Table 2).

### 3’-Tag Sequencing reveals concentration-dependent effects of alcohol on mature oligodendrocytes

Principal component analysis of the top 10,000 most variable genes demonstrated clear separation of control, moderate alcohol, and high alcohol-treated OL groups along PC1 and PC2 (Figure 2A). Gene Set Enrichment Analysis (GSEA) using KEGG, GO, Reactome, Hallmark, and C3 pathways of log fold change between control, moderate, and high alcohol samples identified significant alterations in multiple gene pathways (Figure 2B, Supplemental Table 3). Interestingly, comparisons between the high alcohol and control-treated OL groups revealed a decrease in genes related to ribosomes, cell cycle, and DNA replication.

**Figure 2.**
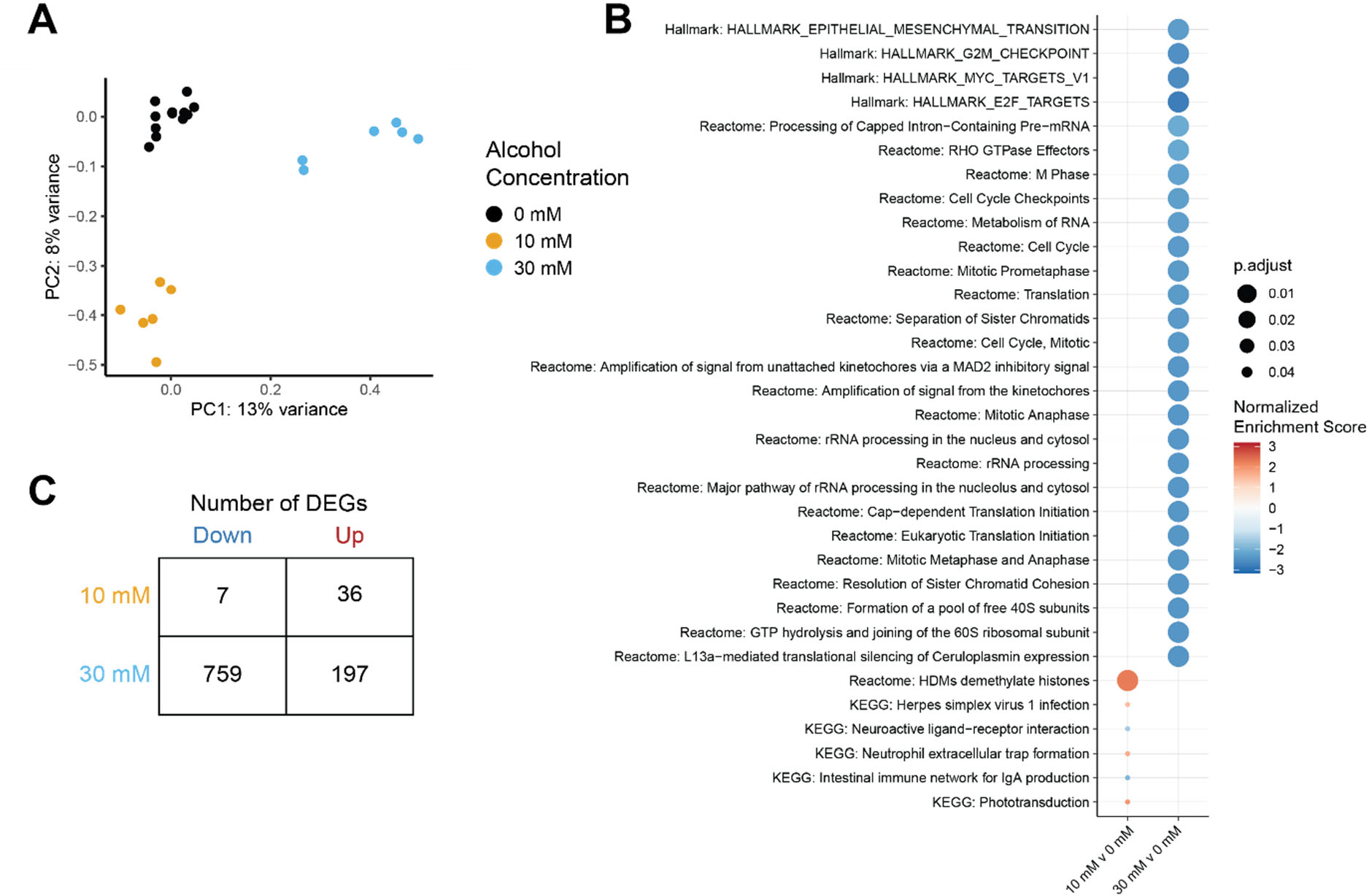
Principal Component analysis and GSEA enrichment testing on concentrationdependent differentially expressed genes. **A)** Control, 10mM, and 30mM alcohol treatment groups separated clearly along PC1 and PC2 in principal component analysis. **B)** GSEA enrichment using KEGG, Hallmark, and Reactome pathways reveals greater pathway perturbations in 30mM alcohol-exposed oligodendrocytes. Adjusted p-values calculated using Benjamini-Hockberg corrections. **C)** Number of up and down DEGs for 10 mM and 30 mM alcohol groups compared to the 0 mM controls.

Differential expression analysis comparing 30 mM and 10 mM to 0 mM controls revealed 956 (197 up and 759 down) and 43 (36 up and 7 down) significant DEGs respectively (defined by p.adjust <= 0.05, Figure 2C, all DEGs reported in Supplemental Table 4). To further identify perturbations in gene expression networks between the high alcohol and control groups, we used Omnipath, a database of known protein-protein interactions, to identify connections between identified DEGs (Figure 3) (Türei, et al., 2021). We observed a significant increase in *MBP* and myelin-associated glycoprotein (*MAG*) in the high alcohol condition. We also observed downregulation of genes in a cluster related to cell cycle regulation centered around cyclin dependent kinase 1 (*CDK1*) and *CDK4*. Other important cell cycle regulators that decreased with high alcohol included Cyclin A (*CCNA2*), Cyclin B (*CCNB1*), and Cyclin D (*CCND1, CCND2*), suggesting that every step of the cell cycle may be affected by high alcohol exposure.

**Figure 3.**
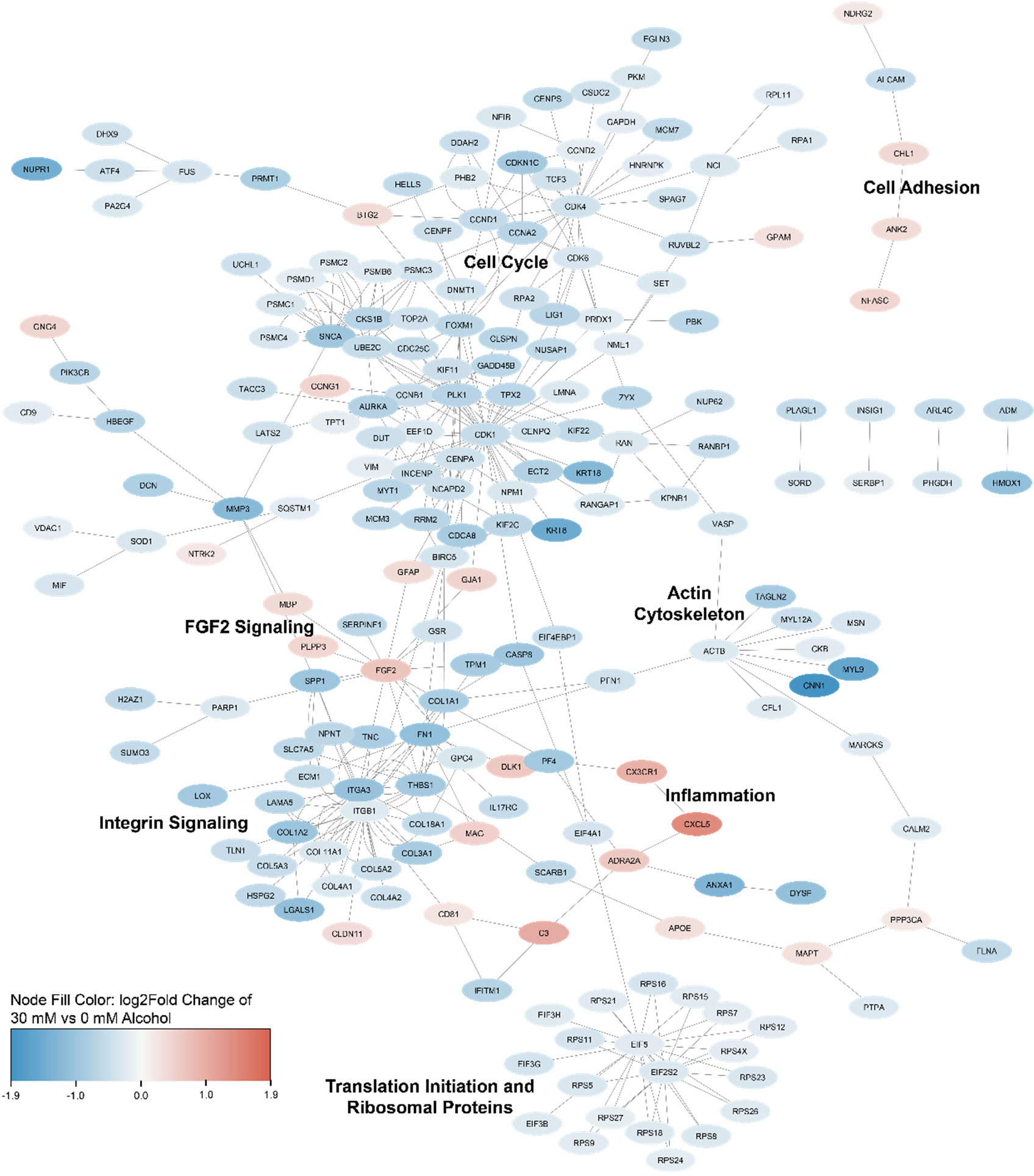
Gene network analysis identifies key changes due to high alcohol exposure. Differentially expressed genes (p.adj <= 0.05) and their known protein-protein interactions from Omnipath were visualized using Cytoscape (line indicates reported interaction in Omnipath). Broad decreases were observed in networks related to protein translation (eukaryotic initiation factors and ribosomal proteins), actin cytoskeleton, integrin signaling, and cell cycle regulation whereas increases were seen in networks related to inflammation and cell adhesion.

On further analysis of cell cycle and myelination genes, we identified additional genes that differed across concentration groups. At the lower alcohol concentration, there were increases in Cyclin B and D and decreases in Cyclin A and E, indicating that low alcohol exposure increased cell cycle progression through G1 and mitosis and decreased progression through G2 and S phase (Figure 4A). At the higher alcohol concentration, all cyclin genes were downregulated. Among known myelin genes (Mayer & Meinl, 2012), we observed increases in *PLP1* and reductions in *PLP2* in both alcohol concentration groups, as well as an increase in MOG expression (Figure 4B). Thus, these data suggest that different concentrations of alcohol may have distinct effects on cell cycle regulation and myelin genes.

**Figure 4.**
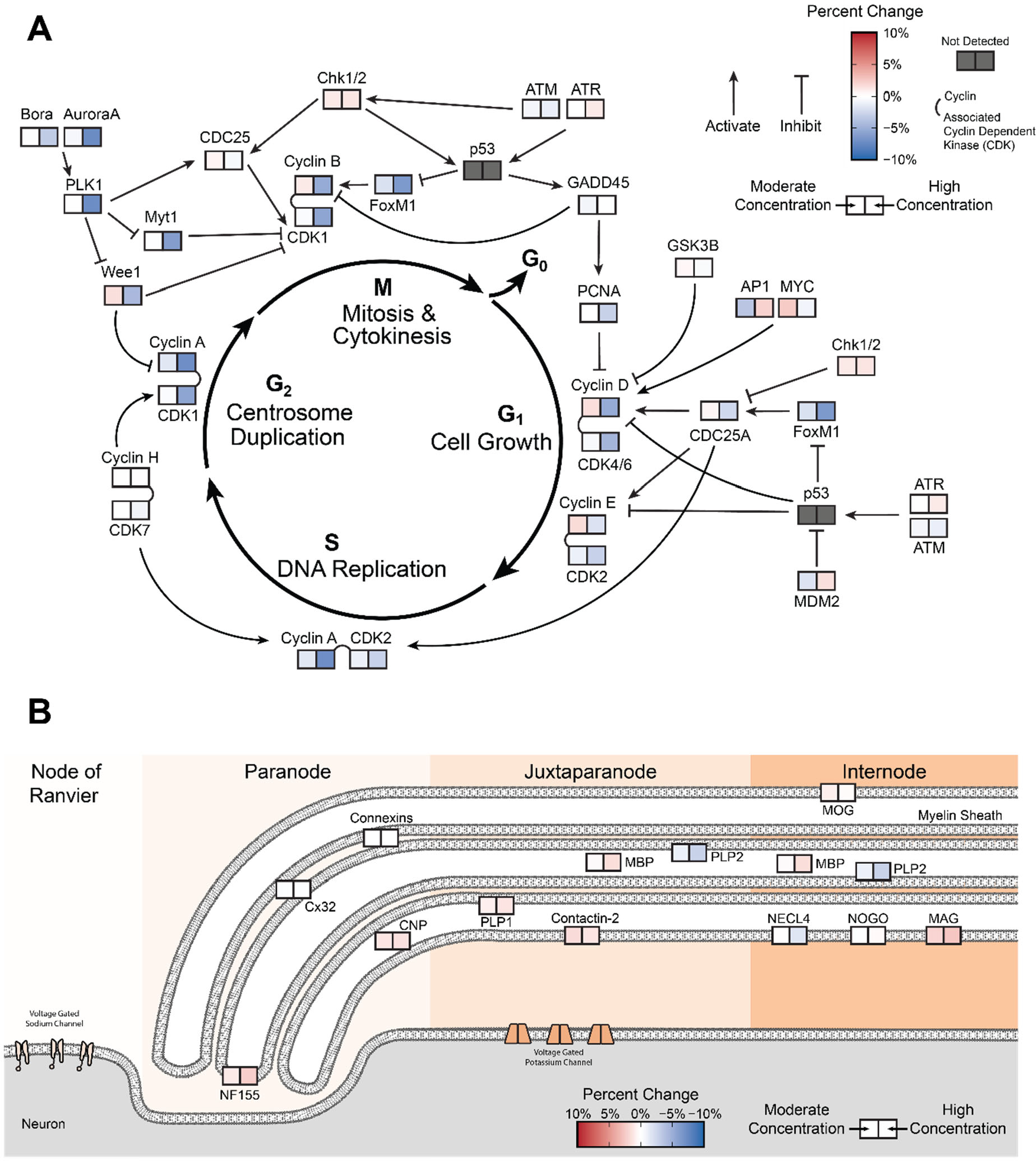
Concentration-dependent changes in cell cycle and myelin gene expression. **A)** Genes involved in cell cycle regulation and progression were analyzed for changes in expression across the moderate and high concentration groups. Figure adapted from GeneTex. **B)** Genes present throughout myelin structures in the internode, juxtaparanode, and paranode regions differed in their expression in the moderate and high alcohol concentration groups. Figure adapted from Mayer and Meinl, 2012.

Another gene cluster in the high alcohol condition revealed broad decreases in eukaryotic initiation factors and ribosomal proteins, suggesting a potential reduction of protein translation in high alcohol-treated cells. Integrin signaling was an additional major cluster, centered on two genes, integrin alpha 3 (*ITGA3*) and beta 1 (*ITGB1*), both of which combine to form α3β1 integrin, which typically interacts with extracellular matrix proteins and is involved in cell signaling. In addition, we observed a decrease in actin cytoskeleton genes and an increase in gene transcript networks involved in cell adhesion, inflammation, and FGF2 signaling.

### Non-linear concentration-dependency of oligodendrocyte gene expression across alcohol treatment groups

To further investigate how alcohol concentration affects gene expression in OLs, we performed pairwise gene comparisons between alcohol conditions and plotted adjusted p values of genes for moderate versus high and control versus moderate alcohol conditions (Figure 5A, Supplemental Table 5). This analysis allowed us to focus on eight unique patterns of significant OL gene expression affected by different alcohol concentrations (represented by eight quadrants in Figure 5A) in comparison to genes that did not change significantly in response to alcohol (middle quadrant, Figure 5A). Genes that were enriched in 10mM (moderate) versus 30mM (high) compared to 10mM versus 0mM (control) conditions were significantly different (p<2.2e-16), indicating that moderate and high alcohol conditions distinctly induced gene expression changes in OLs based on alcohol concentration (Supplemental Figure 1 and Figure 5A). Similarly, ranked-ranked hypergeometric testing confirmed that there was no significant overlap in the transcriptomic changes induced by moderate and high concentrations (p=0.98). Furthermore, when directly comparing log fold change and gene rankings of 10 vs 0 mM and 30 vs 10 mM, the gene profiles were significantly negatively correlated (p<2.2e-16).

**Figure 5.**
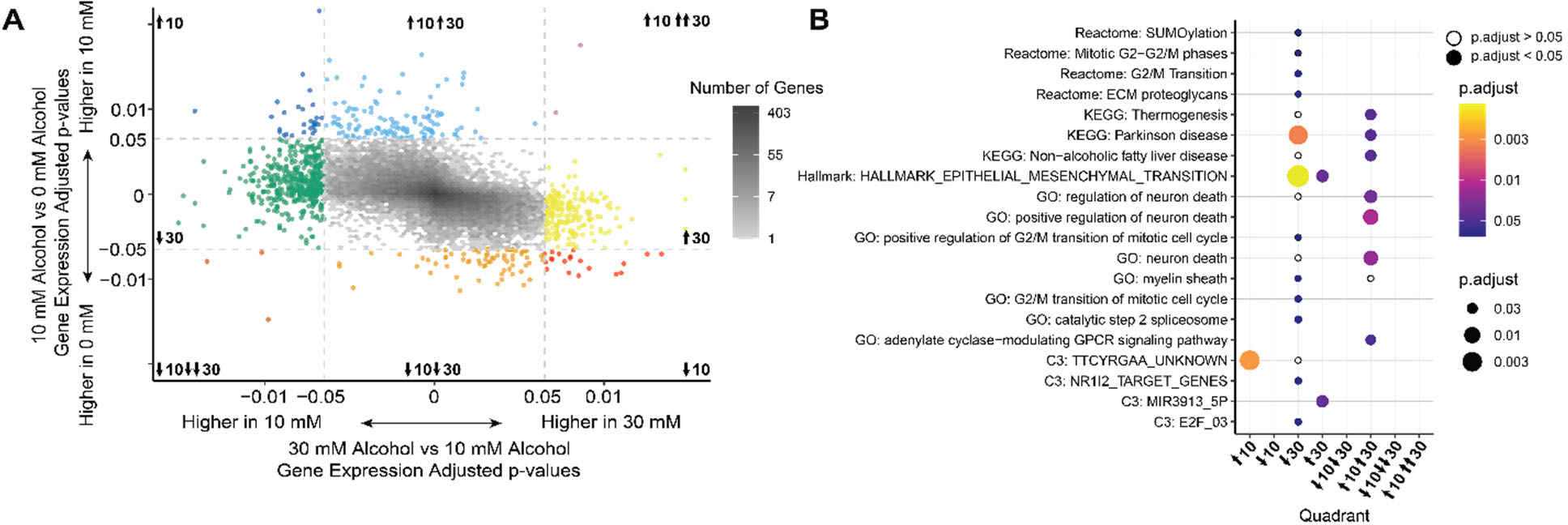
Moderate and high concentrations of alcohol lead to non-linear concentrationdependent changes in gene transcripts. **A)** Pairwise comparisons between alcohol concentrations were evaluated by plotting the significant genes for moderate vs control on the y axis and high vs moderate on the x axis. Genes displayed an overall negative correlation (Pearson’s coefficient = -0.49, p<2.2e-16) across both concentrations’ comparisons, indicating an overall non-linear pattern of concentration dependency on gene expression, with moderate and high alcohol inducing unique genetic changes versus scaling linearly with concentration. **B)** Pathway analysis using Reactome, KEGG, Hallmark, and GO revealed the pathways that were highly represented in each of the quadrants shown in 4A.

Focusing on different quadrants in Figure 5, we demonstrate distinct gene profiles between different alcohol concentration conditions. For example, genes that were significantly higher in the 10mM compared to 0mM and even higher in the 30mM compared to the 10mM (↑10↑↑30 quadrant) represent genes that increase continually with alcohol concentration. Conversely, genes in the ↓10↓↓30 quadrant contained genes that were significantly lower in the 10mM compared to 0mM and even lower in the 30mM compared to the 10mM, indicating that these genes decrease continually with alcohol concentration. The**↑**10**↑**30 quadrant contained genes that had increased expression in the 10mM versus control alcohol comparison and no change in the 30mM versus 10mM alcohol comparison, thus encompassing genes upregulated by alcohol, but non-differentially changed by concentration. Similarly, the ↓10↓30 quadrant contained genes downregulated by alcohol, but non-differentially affected by concentration. The ↓30 quadrant contained genes that had no changes in expression between the 10mM and control groups, but decreased expression in the 30 mM versus 10 mM concentration groups, thus representing genes downregulated only in the high alcohol group (Figure 5A). The **↑**30 quadrant contained genes that had no changes in expression between the 10mM and control groups but increased in the 30mM versus 10 mM comparison group. The ↓10 quadrant contained genes that were only decreased in the 10 mM versus 0 mM concentration groups but increased in the 30 mM versus 10 mM concentration groups, representing genes that were decreased in the moderate group. The **↑**10 quadrant contained genes that were increased in the 10 mM versus 0 mM concentration groups but decreased in the 30 mM versus 10 mM concentration groups.

Evaluation of biological pathways for genes in different quadrants revealed significant changes in gene expression pathways between different alcohol conditions (Figure 5B, Supplemental Table 6). For example, In the ↓30 quadrant, there was a significant reduction in GO terms including actin filament, collagen trimer, extracellular matrix binding, mitochondrial respirasome, mitotic sister chromatid segregation, positive regulation of signal transduction by p53 class mediator, regulation of cell morphogenesis, and SM-like protein family complex. Hallmark’s epithelial mesenchymal transition term was also reduced, in addition to KEGG’s Parkinson’s disease and Reactome’s platelet degranulation. In the **↑**30 quadrant, Hallmark’s epithelial mesenchymal transition term was increased. Furthermore, in the **↑**10**↑**30 quadrant, GO terms including adenylate cyclase-modulating GPCR signaling, neuron death, positive regulation of neuron death, and regulation of neuron death were significantly increased. KEGG’s non-alcoholic fatty liver disease, Parkinson’s disease, and thermogenesis were also increased.

## Discussion

The goal of our study was to better understand how alcohol at different concentrations affects gene expression changes in mature oligodendrocytes (mOL). We found that alcohol was most potent at inducing gene expression changes in mOLs at high concentration, with significant decreases in multiple pathways related to cell cycle regulation, integrin signaling, and protein translation. Additionally, moderate versus high concentrations of alcohol elicited distinct changes in mOL gene expression that differentially impact critical cellular functions, including myelination. These findings support the notion that alcohol can have different biological effects at different concentrations and highlight the critical need for studies to take alcohol dose into consideration.

One interesting finding of our study was that the majority of differentially expressed genes did not change linearly with alcohol concentration. Notably, only two genes, complement 3 *(C3)* and CXC motif chemokine 5 *(CXCL5)* were continually elevated with progressively higher alcohol concentration. *C3* encodes complement 3, the most abundant complement protein, while *CXCL5* encodes an inflammatory cytokine that is a potent stimulator of neutrophil chemotaxis. These key immunological signaling products have been previously observed to increase in systemic alcohol-induced inflammation, and possibly implicate oligodendrocytes in contributing to immune dysregulation via neutrophil recruitment into the CNS (Bykov, et al., 2006, Dominguez et al., 2009, Lowe, et al., 2017). The observed increase of C3 and CXCL5 with alcohol concentration in our study suggests that higher concentrations of alcohol may more potently contribute to neuroinflammation via a neutrophil recruitment mechanism.

Three genes that were continually reduced with progressively higher alcohol concentration included calponin 1 *(CNN1)*, A-kinase anchor protein 12 *(AKAP12)*, and cytochrome P450 1B1 *(CYP1B1). CNN1* and *AKAP12* are associated with cytoskeleton function, whereas *CYP1B1* encodes a member of the cytochrome P450 family of monooxygenases. *AKAP12* is an important scaffold protein that is induced in hepatic stellate cell activation after alcohol exposure (Ramani, et al., 2018). It also plays an important role in OL differentiation in mice, as *AKAP12* deficiency leads to loss of myelin in the corpus callosum (Maki, et al., 2018). The observed decrease in *AKAP12* in OLs at the higher alcohol concentration may point to one potential mechanism for high alcohol-mediated disruption in myelination.

Top KEGG terms that increased as a result of alcohol exposure (without concentration-specific effects) included thermogenesis, Parkinson’s disease, and non-alcoholic fatty liver disease. Top GO terms included pathways related to neuron death and adenylate cyclase-modulating signaling. Notable leading-edge genes in this quadrant included *Mapt*, which encodes Tau protein, *Aplp1*, amyloid beta precursor like protein 1, and *CRHR1*, a GPCR that binds corticotropin releasing hormones. *CRHR1* has been associated with high alcohol intake and in turn, dysregulation of CRH signaling has been linked with stress-induced relapse in animal alcohol studies (Bajo, Cruz, Siggins, Messing, & Roberto, 2008; Treutlein et al., 2006; Ciccocioppo, et al., 2009). Increases in *Mapt* and *Aplp1* as a result of alcohol exposure are interesting findings in the context of Alzheimer’s disease, in which alcohol is known to accelerate cognitive decline (Heymann, et al., 2016).

Pathways that were significantly decreased only in the high alcohol concentration group were KEGG’s Parkinson’s disease and Hallmark’s epithelial mesenchymal transition. Leading edge genes included a member of the importin beta family, Karyopherin Subunit Beta 1 (*Kpnb1)*, which is involved in translocation of cargo across the nuclear pore. The H2 subunit for chromosome condensing complex condensin II, *Ncaph2*, was also in this group, in addition to two proteins involved in protein translation initiation via the 40s ribosomal subunit, Eukaryotic translation initiation factor 3 subunit D *(Eif3d)* and ribosomal protein S5 *(Rps5)*. Significant decreases in *Ncaph2* as a result of alcohol exposure may be indicative of apoptosis, as *Ncaph2* methylation was associated with hippocampal neuron atrophy and cognitive dysfunction in Alzheimer’s disease patients (Shinagawa, et al., 2016). Decreases in *Eif3d* and *Rps5* demonstrate a suppressive effect of alcohol on protein translation. Previous studies have reported that alcohol suppresses protein translation in cardiac muscle (Land, Kimball, Frost, and Vary, 2001), though a similar finding has not yet been reported in oligodendrocytes or other glial cells. Alcohol-mediated suppression of translation may have downstream impacts on myelin production.

Another key gene transcript that was significantly decreased in the high alcohol group was Α3β1 integrin. Integrins are important factors for cell adhesion and interactions between the extracellular matrix and the actin cytoskeleton. Additionally, aberrations in integrin signaling are known to impact OL survival (O’Meara, Michalski, Kothary, 2011). The β1 integrin family of proteins bind to multiple extracellular matrix proteins. Α3β1 integrin in particular, which was one of the key genes in our differentially expressed networks, binds to multiple ligands including fibronectin, collagen IV, nidogen, thrombospondin, laminin-1 and laminin-5 (Belkin & Ann Stepp, 2000). Integrin interactions with laminin are important regulators for OL survival (Decker, Baron, & Ffrench-Constant, 2004). Integrin interactions with axonal laminin-2 also promotes myelin membrane formation (Buttery & Ffrench-Constant, 1999). The observed decreases in integrin gene expression in our study indicates that alcohol exposure to mOLs may lead to decreased mOL survival and myelin production.

In addition to changes in other myelin genes that differed with concentration, we found that high alcohol concentration also significantly increased myelin basic protein *(MBP)* relative to control. MBP is one of the most abundant myelin proteins and is integral to the production and maintenance of myelin. Increase in *MBP* due to alcohol may potentially represent a compensatory mechanism in response to diminished protein translation pathways. Furthermore, the significant increase in *MAG* in the high alcohol group was also notable. MAG is a minor constituent of myelin, but its presence helps stabilize the connection between the myelin sheath and axons (McKerracher and Rosen, 2015). MAG is also a potent inhibitor of neurite growth and can be converted into a soluble form that can affect neighboring cells outside the myelin sheath. The increase in *MAG* due to alcohol exposure may represent a mechanism by which alcohol may inhibit axonal growth following injury (Tang, et al., 1997).

Overall, our results point toward important multimodal effects of alcohol on myelinating genes. Future research should focus on alcohol’s effects in demyelinating diseases such as Multiple Sclerosis (MS), given that MS patients are at a high risk for alcohol use disorder (AUD) compared to the general population (Quesnel & Feinstein, 2004), with up to 40% of individuals exceeding the cutoff for excessive alcohol use on the AUDIT-C (Beier, D’Orio, Spat, Shuman, & Foley, 2014). Additionally, given that most individuals, including MS patients, consume low-moderate amounts of alcohol, future studies need to assess not only high dose but also low-moderate alcohol’s effects on myelination and whether alcohol’s effects may carry a protective or detrimental role on myelination at lower doses.

One limitation of our study was that mRNA alone was studied in these experiments and protein-level analysis would strengthen these findings in future studies. Also, while the *in vitro* model provides important mechanistic information, future studies will need to consider *in vivo* animal or human models to inform how alcohol at different doses may impact demyelination and remyelination.

In conclusion, our study demonstrates that alcohol-induced transcriptomic changes in OLs are concentration-dependent and may have downstream impacts on myelin production. The effects of alcohol on OL demyelination and remyelination could help uncover therapeutic pathways that can be utilized independent of alcohol to aid in remyelinating drug design.

Targeting alcohol-induced changes in cell cycle regulation, integrin signaling, inflammation, or protein translation regulation may uncover mechanisms for modulating myelin production or inhibition. In future studies, it will be important to conduct quantitative protein-based analyses for specific targets related to protein translation. In addition, observations of alcohol’s effects on OL-neuron co-cultures may provide insight into functional aberrations in myelination as a result of alcohol. Further, in vivo studies inducing focal demyelinating lesions in alcohol-consuming animals would be key to assess the effects of dose-dependent alcohol on demyelination and remyelination. Importantly, studying alcohol as a modifiable lifestyle factor could inform patients with chronic demyelinating disorders on the potential risks of alcohol consumption on myelin health. Further research is needed to explore how alcohol consumption affects critical cell types involved in specific demyelinating disorders and AUD in a dose-dependent manner.

## Author Contributions

**Sam A. Bazzi**: Writing – original draft, Writing – review & editing, Conceptualization, Funding acquisition, Formal analysis. **Cole Maguire**: Writing – original draft, Writing – review & editing, Formal analysis. **Dayne Mayfield**: Writing – review & editing. **Esther Melamed**: Writing – review & editing, Conceptualization, Funding acquisition.

## Acknowledgements

We are grateful for the administrative and technical support from Dell Medical School Neurology Department and the UT Austin Biomedical Resource Computational Facility.

---

**Supplemental Figure 1.**
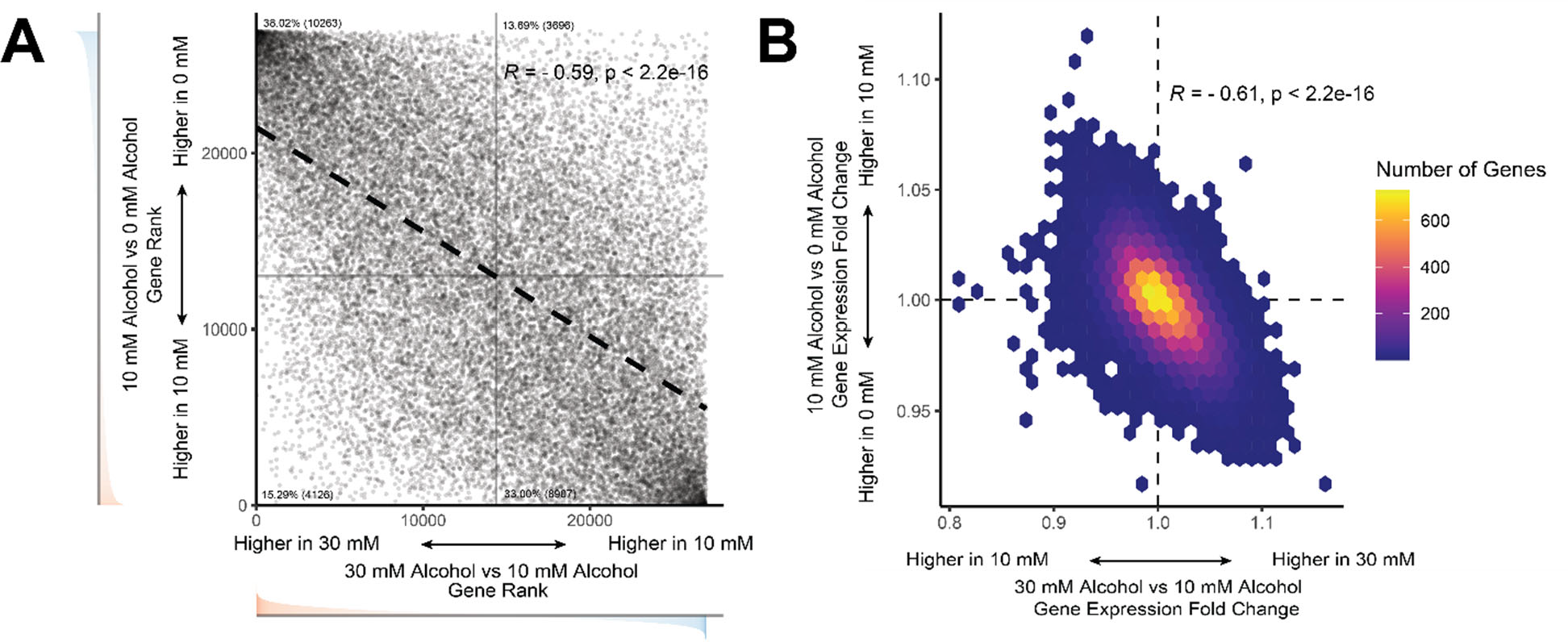
Moderate and high concentrations of alcohol result in concentration-dependent transcriptomics profiles. **A)** Ranking of genes change for 10mM vs 0mM vs 30mM vs 10mM revealed a negative correlation (Spearman correlation R=-0.59, p<2.2e-16) shown as dotted line. **B)** Fold change of 10mM vs 0mM and 30mM vs 10mM also confirmed negative correlation (Spearman correlation R=-0.61, p<2.2e-16).

